# The earliest changes in the translatome upon human T cell activation

**DOI:** 10.1101/2025.07.31.667903

**Authors:** Alexey V. Stepanov, Aoife O’Connell, Phil O’Brien, Patrick B.F. O’Connor, Alla D. Fedorova, Audrey M. Michel, Pavel V. Baranov, Yury P. Rubtsov, Gary Loughran, Dmitry E. Andreev

**Author notes:** contributed equally (Alexey V. Stepanov, Aoife O’Connell).

## Abstract

Stimulation of resting T cells triggers a rapid transition into an activated state, characterized by significant changes in phenotype, metabolism and secretory function. To explore the earliest responses, we examined transcriptional and translational changes in isolated human T cells within the first four hours post-stimulation. Polyclonal stimulation initially upregulates cytokine genes while downregulating certain transcription factors and regulators. Subsequently, selective mRNA translation occurs on 5⍰-terminal oligopyrimidine (TOP) motif-containing mRNAs and *ATF4*, alongside global transcriptional reprogramming involving alternative transcription and splicing. Notably, both stimulated *and* unstimulated T cells undergo dynamic gene expression changes over time, reflecting adaptation to *in vitro* conditions and the loss of *in vivo* homeostatic signals. This includes heightened translation initiation stringency, particularly in non-stimulated cells. Thus, stress responses induced by standard T cell isolation protocols must be considered when investigating T cell signaling and activation.

## Main

Upon stimulation, resting T cells undergo rapid cell growth, followed by clonal expansion 1-2 days later—a process demanding a surge in energy production, protein synthesis, and global reprogramming of the transcriptome and proteome. Resting T cells have low basal protein synthesis rates but are prepared for activation by accumulating a high number of ribosomes and a pool of translationally repressed mRNAs ^1-9^. According to estimations by Wolf and coauthors, resting T cells synthesized in total ∼60,000 proteins per minute, but after 6⍰h of activation, protein synthesis rates increased to ∼300,000 proteins per minute ^6^. The chain of events at the level of gene expression in this early period of activation remains elusive. To address this gap, we employed ribosome profiling ^10^ in human peripheral blood T cells (up to 70 % of these cells are naïve/resting T cells ^11,12^) at 30, 60, and 240 minutes post-stimulation, capturing the immediate dynamics of transcription and translation. Parallel analysis of unstimulated cells distinguished activation-specific responses from stress/adaptation artifacts inherent to *in vitro* culture.

## Results

### Early changes in gene expression in response to stimulation

Human circulating lymphocytes from peripheral blood were used as a source of naïve/resting T cells. In order to minimize the effects of the T cell isolation procedure, we implemented a negative selection of T cells from peripheral blood mononuclear cells of healthy donors. We opted to omit additional separation/purification steps to enrich for particular T cell subsets in order to minimize stressing the T cells. For the purposes of this study, we define total blood T cells as non-activated (resting) T cells ^13^. These cells mostly contain naïve, and various subsets of memory T cells, as well as a small proportion of activated T cells and regulatory T cells.

Half of the isolated total T lymphocytes were stimulated by T cell activator magnetic beads (conjugated with anti-CD3/CD28 monoclonal IgG), while the other half was left without stimulation for the same time points. After 30, 60 and 240 minutes of treatment, cells were pelleted, flash-frozen in liquid nitrogen and subjected to RNAseq and Riboseq (Fig. 1A). Riboseq datasets show strong triplet periodicity and the expected metagene profiles (Extended Data Fig. 1).

**Fig. 1:**
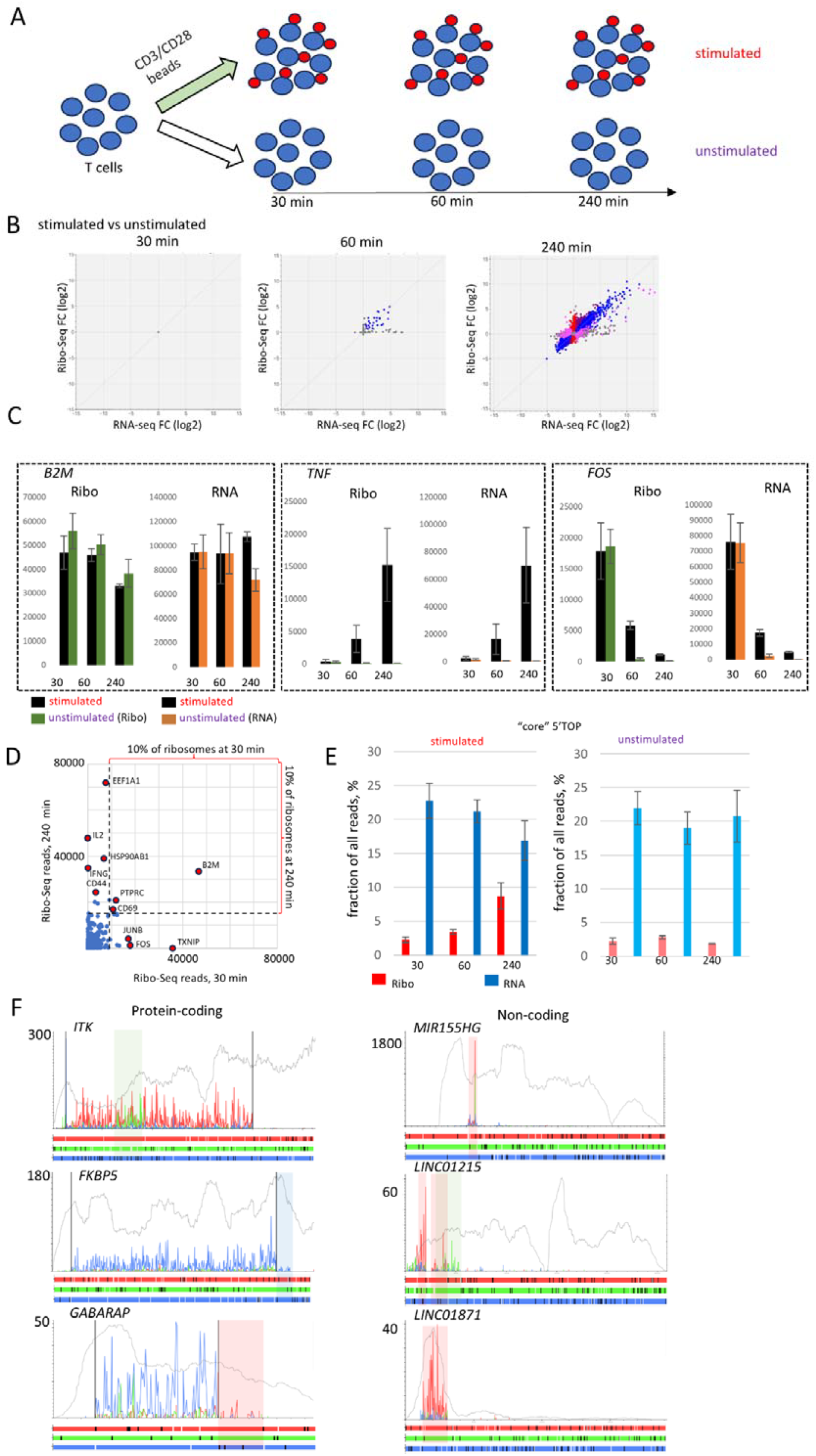
Global changes in transcription and translation during the earliest steps of T cell activation. A) Experimental set-up. B) Differential gene expression of stimulated vs. unstimulated T cells at 30, 60, and 240 minutes. Blue represents DEGs that are regulated at the transcription level with corresponding changes in translation, red - DEGs that are only regulated at the translation level without changes in mRNA abundance, and pink - mixed types of regulation. C) Normalized Riboseq (black and green bars) and RNAseq counts (black and orange bars) for selected genes, with mean and standard deviation error bars from three replicates. D) Riboseq counts for all mRNAs translated at 30 min (x-axis) and at 240 min (y-axis) after stimulation. Selected highly translated mRNAs are highlighted. E) Expression of “core” 5’TOP mRNAs (105 mRNAs) relative to overall expression. Red bars represent the fraction of Riboseq reads from 5’TOP mRNAs compared to all Riboseq reads, with the mean and standard deviation from three replicates. Blue bars represent the same for RNAseq reads. F) Riboseq profiles for selected protein coding and non-coding RNAs in stimulated T cells at 240 min. Riboseq footprints are color coded according to reading frames shown below each plot. RNAseq is shown in grey. Annotated CDS boundaries are shown as black lines while alternatively translated regions are highlighted with different colors.

When comparing the differential gene expression of stimulated versus unstimulated T lymphocytes at the 30 min time-point, not a single differentially expressed gene (DEG) was detected under standard thresholds of statistical significance (Fig. 1B, Table S1). This allows us to use the 30 min time-point as an approximation of the basal gene expression pattern of T cells in the absence of activating signal. At 60 min, 28 mRNAs were upregulated in stimulated T cells at the transcriptional level with corresponding increases in translation (Fig. 1B, Table S2). These include cytokines and chemokines (*TNF, IL13, CCL4, CCL4L2, OSM, XCL1, XCL2*), cytokine ligands and receptors (*IL2RA, TNFSF14, TNFSF9, CD40LG, TNFRSF9*), transcription factors and regulators (*FOS, MYC, KLF6, IRF4, PRDM1, NFKBIA, NFKBIZ*), kinases (*PIM2, MAP3K8*) and genes encoding other immune regulators - *SOCS2, SOCS3, CISH* (suppressor of cytokine signaling family), *ZFP36* (a.k.a. TTP, mRNA binding protein involved in regulation of cytokine mRNA stability ^14^), *IER3* (regulator of cell cycle control and apoptosis ^15^), *GADD45B* (involved in DNA demethylation at specific promoters ^16^) and *GPR171* (G-coupled protein receptor that suppresses T cell proliferation ^17^).

Most but not all of these cases are the result of increased transcriptional output in stimulated T cells. For example, *FOS* mRNA is highly expressed and translated in both stimulated and non-stimulated T cells at 30 min, but at 60 min a relative decrease in mRNA, likely owing to accelerated mRNA degradation, is observed in non-stimulated T cells (Fig. 1C).

At 240 minutes, global gene expression reprogramming is evident between the stimulated and unstimulated states with as much as 4042 DEGs detected (Fig. 1B, Table S3). The repertoire of most highly translated mRNAs (mean Riboseq counts per transcript) drastically changes between 30 and 240 min. At each time-point, out of 14,572 genes that have at least one aligned Riboseq read, the 12 most highly translated genes engage ∼10% of the translating ribosomes. At 30 min they are *B2M, TXNIP, FOS, JUNB, EGR1, CXCR4, DDX5, HLA-B, PTPRC, MYH9, CD69, TMSB4*, and at 240 min - *EEF1A1, IL2, HSP90AB1, IFNG, B2M, CD44, HSPA8, PTPRC, VIM, NCL,CD69, TNF*). Only 3 genes, *B2M, CD69* and PTPRC (*CD45*), are present in both groups (Fig. 1D).

Next, we investigated translational regulation of 5⍰-terminal oligopyrimidine (TOP) motif-containing mRNAs that encode ribosomal proteins and components of the translation machinery. A subset of 105 “core” 5’TOP mRNAs identified previously ^18^, which comprises approximately 23% of all mRNA RNAseq counts in resting T cells, is translated by ∼2% of all ribosomes. After 240 min of stimulation, ∼4 times more ribosomes are engaged in 5’TOP mRNA translation, whereas unstimulated T cells do not induce 5’TOP translation (Fig. 1E).

Given the critical role of protein synthesis in T cell activation, we hypothesized that key translation machinery components would be upregulated to meet heightened biosynthetic demands. Indeed, mRNAs encoding ribosomal proteins and translation elongation factors are activated solely at the translational level, while other components, such as ribosome biogenesis factors and aminoacyl tRNA synthetases, are generally activated at the transcriptional level (Extended Data Fig. 2). For canonical translation initiation factors, the situation is mixed: approximately half of them are activated at the translational level (e.g. *EIF3E*), while the other half are activated at the transcriptional level without any change in translation efficiency (e.g. *EIF3J*). One exception is *EIF1*, which is initially expressed at high levels in resting T cells and does not increase upon activation. Interestingly, *EIF1* mRNA is the most abundant of all translation initiation factors, which may indicate a specific role in T cells. Other notable translationally upregulated genes include *NOP53* and *NSA2*, which are required for 60S subunit biogenesis ^19^. These mRNAs are abundant in resting T cells and their upregulation resembles that of 5’TOP mRNAs. We propose that the translational control of *NOP53* and *NSA2* is essential for activating de novo ribosome biogenesis. *QARS1* (glutaminyl-tRNA synthetase 1) is also upregulated at the translational level, even though its mRNA levels decrease upon activation. In the study by Philippe et al., *QARS1* was found to be regulated by mTOR and LARP1 ^18^. In conclusion, we propose that increasing protein synthesis capacity upon T cell activation requires both translation activation and the execution of transcriptional programs.

Analysis of Riboseq triplet periodicity at 240 min of stimulation allowed us to identify alternative translated regions in protein-coding and putative non-coding RNAs. Some notable examples are shown at Fig. 1F. For *ITK* (IL2 inducible T cell kinase), a nested translon was observed as indicated by an increase of +1 reading frame translation (Fig. 1F, green signal). The nature of this translon is not clear, it may be due to a splicing event (exon skipping) or an alternative transcription start producing a shorter transcript enabling translation initiation at that location. Manual examination of the profile suggests that the translon is at least 84 codons long. In the case of *FKBP5* (FKBP prolyl isomerase 5), Riboseq density downstream of the stop codon points to potential stop codon readthrough resulting in a 29 amino acid C-terminal extension. For *GABARAP* (GABA type A receptor-associated protein), we observed Riboseq signals downstream of the stop codon, but in this case, the ribosomes appear to be translating in a different reading frame (Fig. 1F, red signal). This is unlikely to be +1 ribosomal frameshifting because an increase in ribosome profiling density is observed at the first AUG of the +1 ORF in publicly available Riboseq data (Extended Data Fig. 3), suggesting that this is an independently initiated translon. Since this +1 frame AUG is the first AUG downstream of the CDS start, it is likely reachable by initiating ribosomes due to leaky scanning as observable in other mammalian genes with CDS starts in suboptimal Kozak contexts ^20^. Notably, support for the alternative translons in *ITK* and *FKBP5* can also be observed in publicly available Riboseq data (Extended Data Fig. 3).

In addition, we observed dozens of translons within supposed non-coding RNAs. One interesting example was found in the *MIR155HG* gene, which is the gene for microRNA-155, a well-known master regulator of immune cell responses (reviewed in ^21-23^). Translation of a short translon encoding the peptide *MEMALMVAQTRKGKSVV* was detected (Fig. 1F). Apparently, a fraction of the *MIR155HG* RNA escapes microRNA processing (which occurs in the nucleus) and is instead exported and translated in the cytoplasm. The expression of *MIR155HG* and the translation of the corresponding translon were only detected in stimulated T cells after 240 minutes (Extended Data Fig. 4). Other non-coding RNAs for which translons were observed include *LINC01215* and *LINC01871* (Fig. 1F).

### Stress and adaptation responses associated with cell cultivation conditions

It is expected that T cells isolated from patients experience stress due to their withdrawal from circulation, cell purification, and placement in artificial growth conditions. Therefore, the observed changes in gene expression are a combination of stimulation, response to stress, and adaptation. To uncouple these effects, we examined changes in gene expression in T cells incubated without stimulation over time. A comparison of 30 min vs. 60 min revealed as many as 1,194 DEGs (no stimulation) and 702 DEGs (stimulation), respectively (Fig. 2A, Tables S4 and S5). The number of DEGs increase at 240 min at both conditions (Fig. 2A, Tables S6 and S7).

**Fig. 2.**
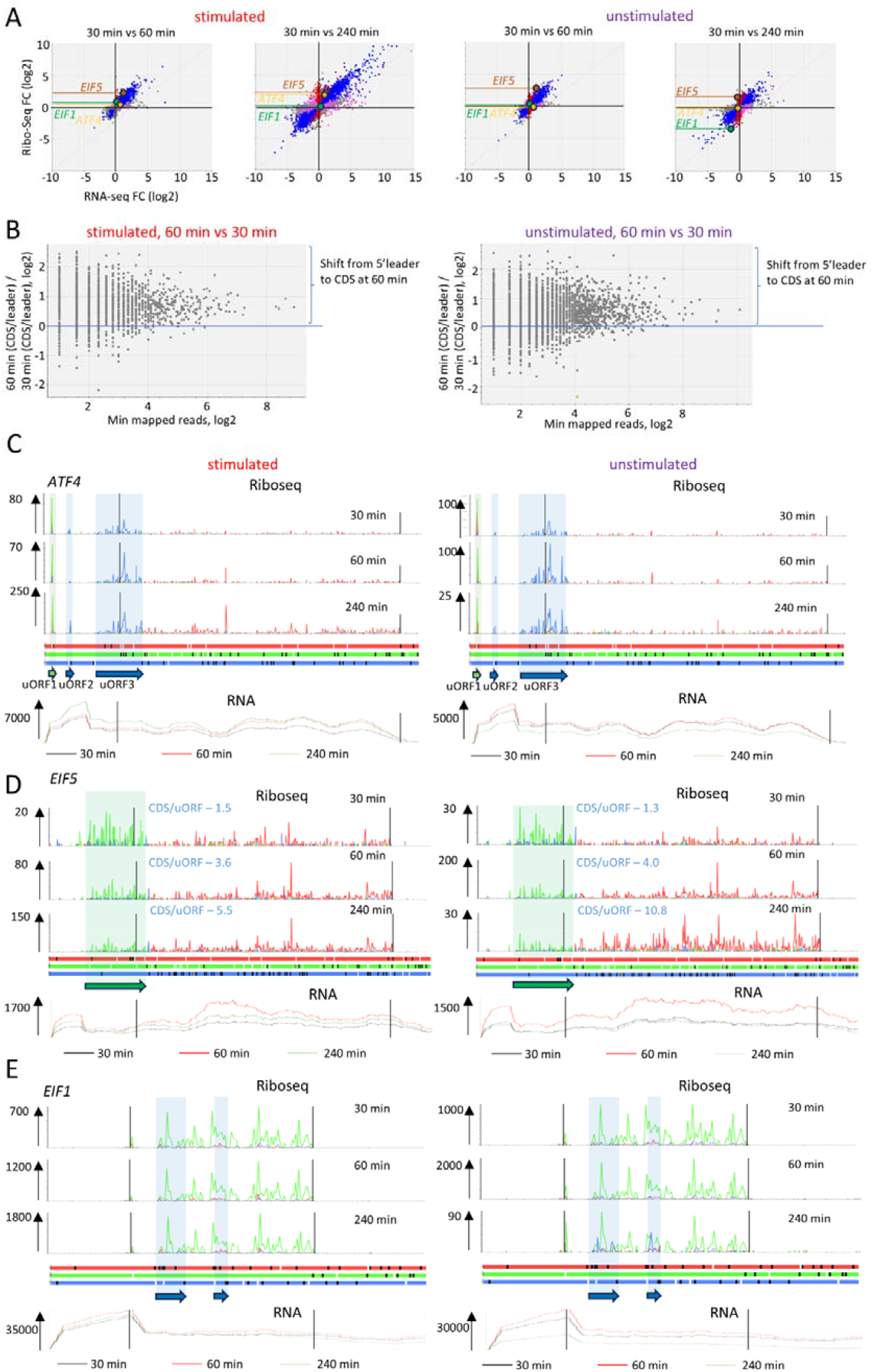
Stress and adaptation responses during T cell cultivation. A) Differential gene expression of stimulated and unstimulated T cells at different time points (60 min vs 30 min and 240 min vs 30 min). The blue dots correspond to genes that are regulated at the transcription and translational level. Red dots represent genes that are only regulated at the translation level, without changes in mRNA abundance. Pink dots show buffered regulation where translation changes compensate for changes in mRNA abundance. The *ATF4, EIF5*, and *EIF1* genes are highlighted. B) Changes in ribosome occupancy at the 5’ leader and CDS for individual genes after 30 and 60 minutes, with and without stimulation. C - E) Riboseq profiles of *ATF4* (*ENST00000674920*), *EIF5* (ENST00000216554) and *EIF1* (ENST00000469257) at different time points with and without stimulation. The Riboseq footprints are color-coded according to the reading frames (provided below diagram). The numbers with arrows indicate the Riboseq scale on the Y-axis. RNAseq data is shown below the profiles. Selected translated open reading frames (ORFs) are highlighted in colored boxes. For *EIF5*, the ratios of CDS/uORF Riboseq reads (aligned to CDS 0 frame (red) and 5’leader +1 frame (green)) are shown in blue.

This indicates that while T cell-related changes in gene expression begin around 1 hour after stimulation, stress- and adaptation-related responses are already at full capacity. This is consistent with the observation that unstimulated T cells undergo significant changes in gene expression (RNAseq) after 6 to 72 hours of cultivation ^24^.

While changes in gene expression mostly occur at the transcriptional level (blue dots in Fig. 2A), a significant fraction of genes are regulated at the translation level. To understand the origin of these translational changes, we calculated the ratios of Riboseq reads in the 5’ leader region to those in the CDS, and compared these ratios between 60 and 30 minutes (Fig. 2B). Both in stimulated and unstimulated cells, the ribosome density shifts strongly from the 5’ leaders to the CDSs, indicating that uORFs may be involved in translational control. We decided to explore known examples of uORF-mediated translation control.

*ATF4* is known to be the main target of translational control regulated during the integrated stress response ^25^. Regulation is mediated by three uORFs and cis-acting elements located within the third overlapping uORF (uORF3) ^26^. At 30 min, ribosome occupancy at uORF1 is high while both uORF2, uORF3 and the CDS have low ribosome occupancies (Fig. 2C).

After 60 minutes, regardless of treatment, the ribosome occupancy at uORF3 increases, indicating translation reprogramming. Interestingly, 240 minutes after stimulation, translation of the ATF4 CDS significantly increases, while unstimulated cells show no change. The activation of ATF4 translation at 240 minutes is accompanied by an increase in uORF2 translation and an increase of the peak at the uORF3 stop codon (Extended Data Fig. 5). While ATF4 translation upregulation is typically linked to stress activation ^27^, we observed it only under stimulated conditions. This suggests that this upregulation may not originate from suboptimal growth conditions (e.g. amino acid deficiency), as both unstimulated and stimulated cells grow in the same medium. One can hypothesize that when T-cell activation rapidly increases global translation, it depletes either the eIF2*GTP*tRNAi (TC) pool or the pool of available ribosomes, which in turn drives ATF4 induction without the involvement of eIF2 phosphorylation, similar to the recently described non-canonical integrated stress response pathway ^28^.

Another example is the *EIF5* mRNA, which contains an overlapping uORF with three AUG codons in a poor Kozak context. It has been demonstrated that eIF5 regulates the translation of its own mRNA, with increased levels of eIF5 enhancing uORF translation and inhibiting CDS translation ^29^. At 60 minutes, regardless of treatment, the ribosome density dramatically shifts from the uORF towards the CDS. After 240 minutes, the translation of the uORF is further inhibited, but in unstimulated cells, the shift from the uORF to the CDS becomes more pronounced (Fig. 2D, right panel). This suggests that the stringency of start codon selection may increase during incubation, especially in the absence of stimulation.

If this is the case, it is expected that *EIF1* translation may also be regulated. *EIF1* mRNA has an AUG start codon in a weak Kozak context, and a high level of eIF1 inhibits AUG recognition and its own translation ^30^. No significant changes in Riboseq profiles for *EIF1* were observed under stimulation conditions, but after 240 minutes without stimulation, there were drastic changes. Translation of *EIF1* was strongly downregulated at this point, and this was accompanied by the activation of translation of two short nested translons (*i*.*e*. within the CDS). This suggests that the first AUG was skipped due to leaky scanning, and initiation occurred at downstream AUG codons (Fig. 2E). *EIF1* mRNA levels also decrease, which we speculate may be because the translation of the nested translons destabilizes the mRNA.

Interestingly, *EIF1B*, a paralog of *EIF1*, is also similarly regulated at the translation and transcription levels: at 240 minutes without stimulation, the translation of *EIF1B* decreases, accompanied by translation of short translon located in the 3’ trailer (Extended Data Fig. 6). We also investigated the translation of *BZW1* mRNA, which has been reported to be regulated by stringency ^31^. Similar to *EIF1, BZW1* has an evolutionarily conserved poor AUG initiation context. At 240 minutes of incubation, the translation of two downstream short translons, likely originating from increased leaky scanning of the *BZW1* poor start codon ^20^, is activated, and this effect is more pronounced in unstimulated cells (Extended Data Fig. 6).

Collectively, these findings suggest that both stimulated and unstimulated cells experience a decrease in initiation at non-optimal start codons (increased stringency). However, this phenomenon is more pronounced in unstimulated cells. Therefore, increased translation initiation stringency may be considered a stress/adaptation response. Activation stimuli, on the other hand, counteract the shift in stringency to provide an optimal translation environment for the activated state.

Some translation regulatory events are identical in both stimulated and unstimulated cells, such as efficient activation of *FTL* mRNA translation (Extended Data Fig. 6). *FTL* is regulated at the translation level through an iron responsive element (IRE) located in its 5’ leader region in response to changes in intracellular iron levels ^32-34^. This regulation may indicate iron deficiency in the cell culture medium.

### Examples of novel gene specific regulation events that may be important for T cell responses

Changes in Riboseq profiles (silhouettes) on a particular gene locus between different conditions may occur because of changes in the relative translation of translons and/or because of changes in the composition of mRNA isoforms expressed from that locus. We compared Riboseq profiles for individual mRNA isoforms at different time points to uncover numerous regulatory events. Some of these events, to the best of our knowledge, have not been previously described in relation to T cell physiology. We provide several examples of these regulatory events, which occur either exclusively in stimulated cells (stimulation-specific events) or in both stimulated and unstimulated cells (adaptation/stress response-specific events).

*TNFSF8* (a.k.a. *CD153* or *CD30L*) encodes a protein that belongs to the tumor necrosis factor (TNF) ligand family. Its cognate receptor is CD30. *TNFSF8* plays a role in role in controlling mycobacterial infections causing tuberculosis ^35^ and leprosy ^36^, as well as regulating senescence-associated (SA) CD4^+^ T cell activity ^37^. When CD153 is engaged on SA-T cells after TCR stimulation, it associates with the TCR/CD3 complex, restoring TCR signaling and promoting T cell activation ^37^. We observed that at 30 minutes in both stimulated and unstimulated T cells, translation of the *TNFSF8* CDS is likely suppressed by the translation of a uORF (Fig. 3A). Upon stimulation, translation of the CDS greatly increases, and translation of the uORF disappears. Examination of the RNAseq profile reveals an alteration in the silhouette. Analysis of the genomic alignment reveals that, upon T cell stimulation, a novel transcription start site (TSS) is activated. This TSS not only removes the entire uORF, thereby derepressing translation, but it also removes the first AUG codon of the *TNFSF8* CDS, resulting in translation of a N-terminally truncated protein (Fig. 3A). The 11 amino acids absent from the novel proteoform (MDPGLQQALNG) are located in the cytoplasmic tail of CD153. Therefore, such a deletion could affect signaling associated with the interaction between transmembrane receptor CD30 (TNFRSF8) and its ligand CD30L (CD153, TNFSF8).

**Fig. 3:**
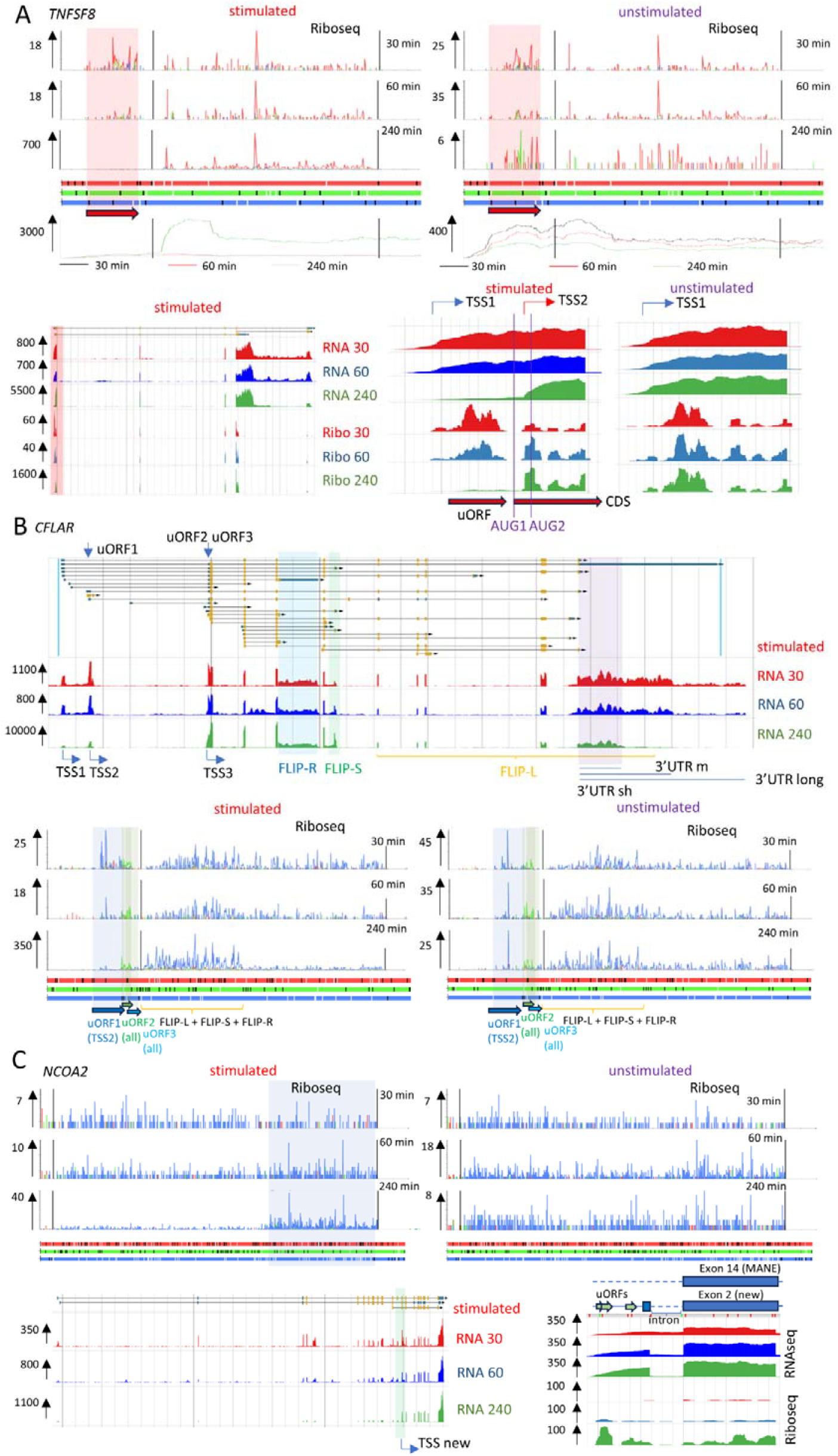
Alternative transcription starts usage during T cell stimulation. A) Riboseq profiles for *TNFSF8* (*ENST00000223795*) with and without stimulation. Riboseq footprints are color-coded according to the reading frame bias to color-match the supported frame in the ORF plot below. The numbers with arrows indicate the Riboseq scale on the y-axis. Translon locations are differentially highlighted. The genomic alignment of Riboseq and RNAseq for *TNFSF8* is provided below the profiles. B) The genomic alignment of RNAseq for the *CFLAR* locus. Regions of interest are highlighted with arrows and color shading. Riboseq profiles for *CFLAR* (*ENST00000423241*) are displayed below. uORFs are highlighted. The numbers with arrows indicate the Riboseq scale on the y-axis. C) Riboseq profiles of *NCOA2* (*ENST00000452400*) with and without stimulation. The Riboseq footprints are color-coded to match supported reading frames in the ORF plot below. The numbers with arrows indicate the Riboseq scale on the y-axis. The region where Riboseq signal density changes most dramatically are shaded in blue. The genomic alignment of Riboseq and RNAseq for *NCOA2* is provided below the profiles.

*CFLAR*, also known as c-*FLIP*, encodes a CASP-8 and FADD-like apoptosis regulator that is a master regulator of apoptosis and immune signaling in various normal and cancer cells (see recent review in ^38^). The complex organization and regulation of the *CFLAR* locus allows for the expression of many transcript isoforms, including three major ones that encode three proteins: long c-FLIP_L, and two shorter variants c-FLIP_S and c-FLIP_R. Each of these proteins contains different combinations of death effector and caspase-like domains. The ratio of these isoforms can influence the cellular decision to undergo apoptosis or other forms of programmed cell death, such as necroptosis ^39^. Before stimulation, T cells express all three mRNA transcripts that encode c-FLIP_L, c-FLIP_S, and c-FLIP_R (Fig. 3B). The most abundant transcript is the longest, c-FLIP_L. At 240 minutes post-stimulation, this pattern shifts towards transcripts encoding c-FLIP_S and c-FLIP_R. For c-FLIP_L, the 3’UTR is significantly shortened. Before stimulation, two major transcription start sites, TSS1 and TSS2, were used. At 240 min post stimulation, TSS3 is activated and predominates over TSS1 and TSS2. As a result, three different 5’leaders exist, all 5’leaders contain two uORFs (17 triplet long uORF2 and 25 triplet-long uORF3) located close to the CDS start codon, while 5’leader 2 (which originates from TSS2 usage) contains an additional 58 triplet-long uORF1. Without stimulation, uORF1 is translated, likely repressing translation of the corresponding CDS. Stimulation-induced switching to TSS3 removes uORF2, so translation of FLIP isoforms may be derepressed. Therefore, T cell stimulation-dependent changes in *CFLAR* expression are also regulated by the selection of alternative 5’leaders with different uORF composition.

*NCOA2* (a.k.a. *SRC2*) encodes for a protein that belongs to the p160 SRC family of nuclear receptor coactivators involved in various cellular processes, including T cell differentiation and immune responses ^40^. In naïve T cells, NCOA2 plays a role in promoting Treg cell differentiation ^41^. Mechanistically, NCOA2 is recruited by NFAT1 to the promoter of NR4A2, activating its expression and subsequently stimulating FOXP3 expression. In addition, NCOA2 promotes CD8+ T cell-mediated immune responses by stimulating T-cell activation via upregulation of PGC-1α ^42^. We observed an increase in the translation of the C-terminal fragment of NCOA2 in response to stimulation (Fig. 3C). Analysis of genomic alignments suggests that in stimulated T cells, a novel transcription start site located upstream of exon 14 of the *NCOA2* (the principal MANE isoform) is activated, giving rise to a previously unannotated, shorter mRNA transcript. This new transcript joins the 5’ terminal exon, which contains several uORFs and an AUG start codon that is in frame with the CDS, with exon 14, creating a shorter CDS. The novel NCOA2 protein form should clearly have a functionally distinct role from the known long NCOA2 form, as it lacks the bHLH, PAS, and RID domains which are important for signal transduction and binding of coregulators ^43^.

We also observed changes in Riboseq profiles over time during T cell incubation that are not specific to stimulation. One example is the case of *CCNT1* (Cyclin T1). Cyclin T1 forms a complex with CDK9 ^44^, which is known as positive transcription elongation factor b (p-TEFb). We observed that at 30 minutes of T cell incubation, Riboseq density drops on the mRNA in the region of the exon 6 – exon 7 boundary (Fig. 4A). Genomic alignment showed that at early time points, exon 7 was skipped, resulting in translation termination within exon 8. However, over time, exon 7 became included, leading to full-length CCNT1 synthesis. Urano et.al. ^45^ discovered a *CCNT1* splice variant without Exon 7 (deltaE7) and showed that the deltaE7 transcript is abundant in non-proliferating cells and tissues. Experiments using constructs encoding the CCTNT1 deltaE7 protein demonstrated its physical interaction with CDK9 and competition with full-length CCNT1 for binding to CDK9.

Similar changes were observed for the *OGT* gene, which encodes for O-GlcNAc transferase (Fig. 4B). Similar to *CCNT1*, at 30 minutes after T cell incubation, the Riboseq density dropped at the mRNA region near the exon 3 - exon 4 boundary. The expression of *OGT* has been reported to be regulated by detention of a highly conserved intron (DI), in response to the inhibition of OGT activity ^46^. We observed a rapid increase in DI splicing over the course of incubation, regardless of whether the cells were stimulated or not. At 30 minutes, when the DI was not spliced, we noticed the presence of Riboseq density within the DI, corresponding to a stop codon in the first reading frame. This suggests that DI-containing *OGT* mRNA has been exported to the cytoplasm and is undergoing translation. It also suggests the possibility of translation of a novel truncated OGT proteoform, similar to that of CCNT1, which exists during the quiescent state.

*SRP19* codes for a subunit of the signal recognition particle (SRP). It is the first subunit to bind to 7SL RNA during SRP assembly, and it is rate-limiting for SRP maturation ^47^. Analysis of Riboseq profiles and genomic alignments (Fig. 4C) revealed that after 30 minutes of incubation, T cells express and translate two mRNAs with alternative 3’ terminal exons, resulting in the synthesis of SRP19 and an SRP19 variant with an altered C-terminus. While the functional role of this novel variant is not yet clear, it is possible that it may not be able to engage in SRP, limiting SRP maturation ^48^. Over time, T cells begin to express the conventional *SRP19*, which should maintain their secretory capacity.

**Fig. 4.**
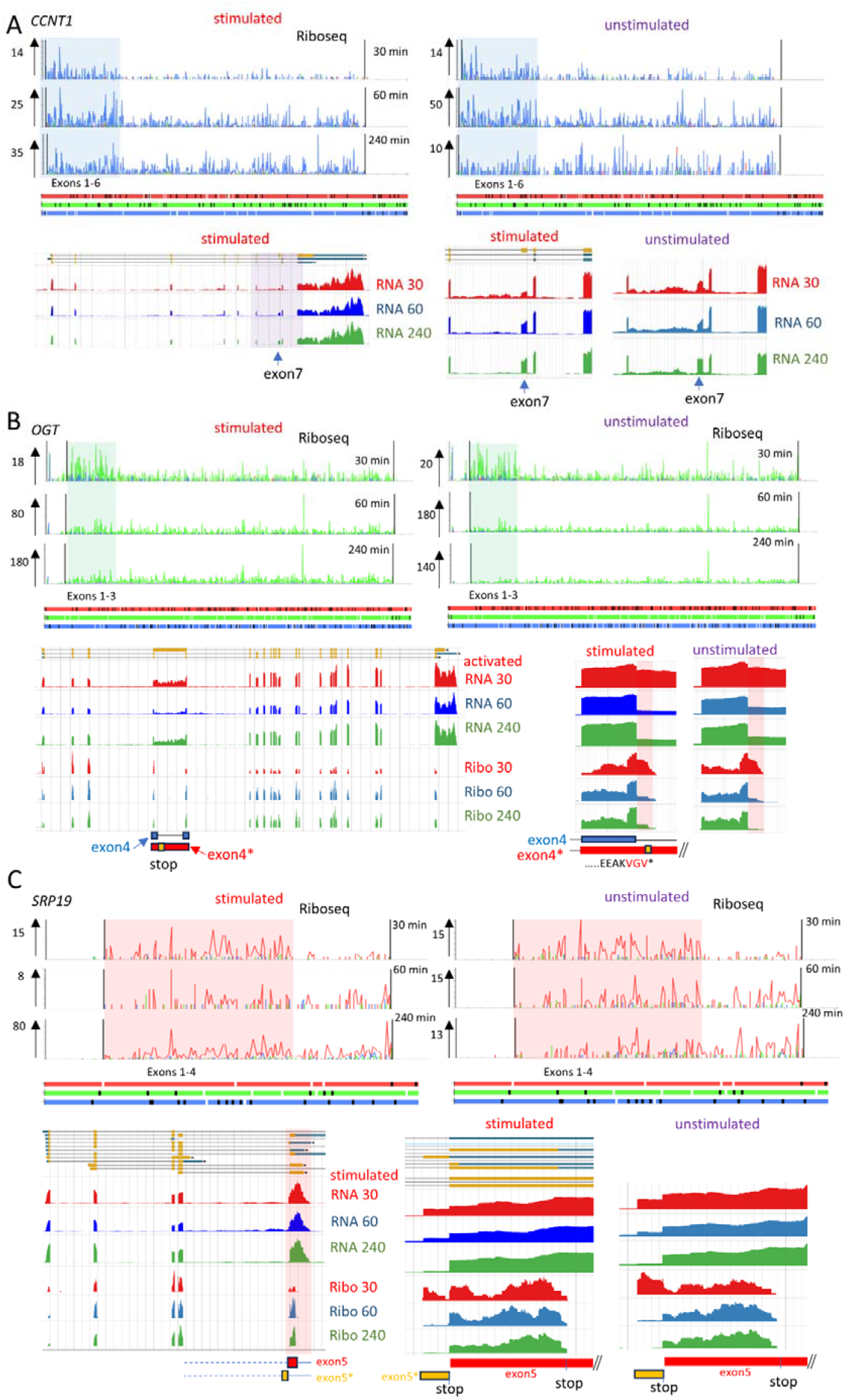
Alternative splicing events in response to T cell cultivation. A) Riboseq profiles for *CCNT1* (*ENST00000261900*) with and without stimulation. Riboseq footprints are color-coded to match supported reading frames in ORF plot below. The numbers with arrows indicate the Riboseq scale on the y-axis. The region where Riboseq density changes is highlighted. The genomic alignment of Riboseq and RNAseq for *CCNT1* is provided below the profiles. B) Riboseq profiles for *OGT* (*ENST00000373719*) with and without stimulation. The genomic alignment of Riboseq and RNAseq for *OGT* is provided below the profiles. C) Riboseq profiles for *SRP19* (*ENST00000505459*) with and without stimulation. The genomic alignment of Riboseq and RNAseq for *SRP19* is provided below the profiles.

## Discussion

Resting/naïve T lymphocytes maintain viability through selective mRNA translation while stockpiling translationally repressed mRNAs and excess translation machinery required for future activation. Ribosome profiling has allowed us to understand the general principles of translation control in T cells in a quiescent state and after receiving activation stimuli. Stimulation for 240 min revealed 521 genes with significant alterations in CDS ribosome occupancy, with 70% (363 genes) showing increased translation (Table S8). In contrast, incubation without stimulation yielded 339 differently translated genes, with 49% (167 genes) showing increased translation. Notably, just 54 mRNAs showed overlapping translational regulation between the two groups, demonstrating that T cell activation initiates a distinct translational program that overrides baseline stress/adaption responses. These findings establish that most instances of translation activation occur upon stimulation.

To demonstrate the core translatome dynamics in T cells, we visualized the most abundant mRNAs at the translation level (Figure 5). In resting T cells, the 200 most abundant mRNAs (∼48% of all RNA sequences aligned to coding sequences, excluding mitochondrial mRNAs) account for approximately 25% of all ribosomes involved in translation (Table S9). For these mRNAs we calculated ribosome occupancy (also frequently referred as translation efficiency (TE), although this term is less accurate) as a ratio of Riboseq to RNAseq within their CDSs under different conditions. Out of 200 mRNAs, approximately half had low ribosome occupancy prior to stimulation (stimulation for 30 minutes), which increased after 240 minutes of stimulation, but did not change after incubation without stimulation (240 minutes unstimulated). These abundant mRNAs, which are translationally repressed in resting T lymphocytes, represent known 5’TOP mRNAs and some novel genes, which, to our knowledge, were not reported to contain functional 5’TOP motifs. These include 60S ribosome biogenesis factor *NOP53* and cell surface receptor *CD48* (Table S9). In the group with low ribosome occupancy, only 3 mRNAs, *MLLT6, PPP1R15A* and *UBC*, were not translationally de-repressed upon stimulation, indicating a different mechanism of translation repression. For instance, translation of *PPP1R15A*, which encodes a regulatory phosphatase subunit involved in dephosphorylation of eIF2, is controlled by two uORFs in a stress dependent manner ^49-51^.

**Fig. 5:**
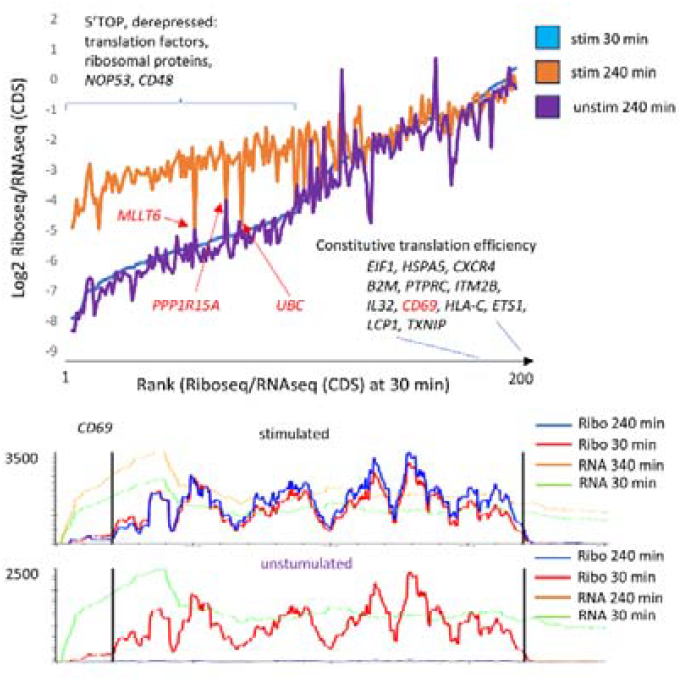
General overview of translation response in T cells. Changes in ribosome occupancy for the 200 most abundant mRNAs. Selected genes that are not derepressed upon stimulation are marked with red arrows. The Riboseq and RNAseq profiles for the *CD69* mRNA (ENST00000228434) under different conditions are shown below.

Another group of abundant mRNAs, which are located on the right side of the graph (Figure 5), have a high ribosome occupancy in resting T cells, which does not change over time under both stimulated and unstimulated conditions. This suggests that de-repression of selected mRNAs, rather than a general activation of translation, is the main source of global changes in the translatome. Therefore, after 4 hours of stimulation, there are three main pools of translated mRNAs: 1) de-repressed mRNAs with 5’TOP motifs (e.g., *EEF1A1*), 2) constitutively translated mRNAs (e.g., *B2M*), and 3) newly transcribed mRNAs (e.g., IL2). Comparing the results of SILAC labelling and RNA sequencing, Wolf et al. reported that in naïve CD4+ T cells, there is a pool of 242 mRNAs that are translationally repressed ^6^. While the list of translationally repressed mRNAs overlaps largely with those identified in our study, there are some important genes with different expression patterns. For example, the *CD69* mRNA, which was reported to be translationally repressed, is efficiently expressed and translated in our experiments at 30-240 minutes of stimulation, without changes in ribosome occupancy. However, without stimulation, the *CD69* mRNA levels dramatically decreases (Fig. 5). The most likely explanation for this discrepancy is that the isolation of naïve CD4+ lymphocytes and labelling with heavy amino acids takes a significant amount of time. During this time, before stimulation, gene expression changes occur. Therefore, it is important to consider that non-stimulated T lymphocytes undergo significant changes in gene expression during the cultivation. A significant limitation of this study was that a mixed population of T lymphocytes, rather than individual subtypes, was investigated. Therefore, some changes in gene expression may occur in specific cell subtypes that we currently cannot identify. Development of new protocols for rapid and nontraumatic isolation of specific T-lymphocyte types will allow researchers to address this issue in the future.

## Methods

### T cell isolation and activation

Human peripheral blood mononuclear cells (PBMCs) were isolated from whole blood obtained from five healthy donors through Ficoll-Paque (Millipore) density gradient centrifugation. Samples were collected with the approval of The Scripps Research Institute’s Institutional Review Board and in accordance with ethical guidelines. T cells were purified from PBMCs using the EasySep Human T Cell Isolation Kit (STEMCELL Technologies).

For each donor, the isolated T cells were divided into six aliquots. Three of these aliquots were activated with CD3/CD28 beads at a 1:1 bead-to-cell ratio (ThermoFisher Scientific) in Aim V medium (ThermoFisher Scientific) supplemented with 40 IU/ml recombinant IL-2 (R&D Systems). Activation was carried out for 240 minutes. Untouched T cells were maintained in Aim V medium without activation and cytokine stimulation.

To arrest eukaryotic translational elongation for ribosome profiling, cells were harvested at 30, 60, and 240 minutes post-stimulation. At each time point, cells were diluted tenfold with ice-cold PBS containing cycloheximide at 1 μg/ml to halt translation elongation. Cells were then pelleted by centrifugation and snap-frozen for downstream processing.

### Ribosome Profiling

Ribosome profiling was adapted from ^52,53^ with modifications to the procedure. Human T-cell pellets were placed on ice and resuspended in 250µl polysome lysis buffer (20 mM Tris–Cl (pH 7.5), 150 mM NaCl, 5 mM MgCl2, 1 mM DTT, 1% Triton X-100, 0.1 mg/ml cycloheximide, 25 U/ml TURBO DNase (Invitrogen), in nuclease-free water). Cell lysates were incubated on ice for 10 mins and then centrifuged at 10,000 × g for 5 mins at 4°C to pellet cell debris. RNA concentration of the supernatant was measured using the Qubit 4.0 Fluorometer and Qubit RNA Broad Range (BR) Assay Kit (Invitrogen). Aliquots were taken for Riboseq (∼3-7 µg) and RNAseq (∼1 µg) library preparation.

For RNAseq, total RNA was extracted from lysate using TRIzol Reagent (Invitrogen) followed by clean-up using the RNA Clean and Concentrator-5 Kit (Zymo Research), both according to manufacturers’ instructions. The RNA concentrations and RNA Integrity Number were measured using the 2100 Bioanalyzer System and the RNA 6000 Nano Assay Kit (Agilent). The RNAseq libraries were then generated at GENEWIZ (Azenta Life Sciences, Germany).

For Riboseq, lysates were treated with 20U RNase I (10 U/µl, Biosearch Technologies) at 23°C for 45 mins at 400 rpm. 80U SUPERaseIn RNase Inhibitor (20 U/μL, Invitrogen) was added to quench the digestion. 10µl 10% SDS was then added and the RPFs were then purified using the RNA Clean and Concentrator-5 Kit according to manufacturer’s instructions. The RNA was dephosphorylated by treatment with 10U T4 PNK enzyme (New England BioLabs) in a 10μl reaction supplemented with 10X PNK buffer and 20U of SUPERaseIn RNase Inhibitor for 1 hour at 37°C, and then purified using the Oligo Clean and Concentrator Kit (Zymo Research) according to manufacturer’s instructions. RNA was resolved by 15% denaturing polyacrylamide TBE-urea PAGE (containing 1X TBE, 8 M urea, 40% acrylamide (19):bis-acrylamide (1)). Bands corresponding to RNA fragments of 28–32 nt were excised and extracted using the ZR small-RNA PAGE Recovery Kit (Zymo Research) according to manufacturer’s instructions. The 3’ ends of the RNA were then ligated with a pre-adenylated DNA linker with a 3 nt unique molecular identifier (purchased from IDT, 5’-/5Phos/(N:25252525)(N)(N)AGATCGGAAGAGCACACGTCTGA/3ddc/) for 3 hours at 23°C. The 15.5µl reaction was supplemented with 1 µM pre-adenylated linker, 17.5% PEG-8000, 1X T4 RNA Ligase Buffer, 100U T4 RNA Ligase 2, truncated K227Q (200U/μl, New England BioLabs). The ligation reactions were separated by 15% denaturing polyacrylamide TBE-urea PAGE as previously described and the ligation products were excised and extracted using the ZR small-RNA PAGE Recovery Kit. This RNA-DNA hybrid molecule is used as a template for reverse transcription.

cDNA synthesis and PCR of 3’ ligated products were carried out using the D-Plex Small RNAseq Kit for Illumina with Unique Dual Indexes (Diagenode) with the following modifications to the procedure. Step 2: RNA Tailing was omitted and in Step 3: Reverse Transcription with Template Switching: Step 3.1. 0.1 nM of a custom reverse transcription primer (purchased from IDT, 5’-GTGACTGGAGTTCAGACGTGTGCTC-3’) was added instead of the Reverse Transcription Primer (RTP) provided. The remaining procedures for Reverse Transcription and PCR were carried out according to the manufacturer’s instructions. The PCR product was purified using the Select-a-Size DNA Clean & Concentrator Kit (Zymo Research) using the ≥150bp (70µl) protocol, according to manufacturer’s instructions. Libraries were pooled for shallow sequencing on the Illumina iSeq 100 System.

### rRNA Depletion of Riboseq Libraries

For rRNA depletion, libraries were pooled and subjected to CRISPR-Cas9 cleavage using a single-guide RNA (sgRNA) pool designed from the most-abundant contaminating rRNA-derived sequences, including a PAM sequence, adapted from (32345633). The sequences were purchased from IDT as ssDNA oligonucleotide templates which underwent anneal and fill-in reactions to form dsDNA templates for in-vitro transcription. Using separate reactions, 5X Phusion HF Buffer (NEB) was added to 10 µM of each ssDNA oligo and 10 µM of a scaffold anti-sense primer (purchased from IDT, 5’-AAAAGCACCGACTCGGTGCCACTTTTTCAAGTTGATAACGGACTAGCCTTATTTTAACTTGCTATTTCTAGCTCTAAAAC-3’) and incubation consisted of denaturation at 95°C for 5 min, annealing from 95°C to 50°C at 0.1°C/sec, hold at 50°C for 5 mins, then 50°C to 37°C at 0.1°C/sec. Following annealing, 10 mM dNTPs (NEB), 2U Phusion HF DNA Polymerase (NEB) and nuclease-free water were added to a final volume of 50µl. The fill-in reaction consisted of denaturation at 98°C for 30 secs, 5 cycles of 95°C for 10 secs, 50°C for 15 secs, 72°C for 10 secs, and a final extension at 72°C for 5 mins. The dsDNA was purified using the Select-a-Size DNA Clean & Concentrator Kit using the ≥50bp (300µl) protocol, according to manufacturer’s instructions. In-vitro transcription of the dsDNA was performed using T7 RiboMAX Express Large Scale RNA Production System (Promega) according to manufacturer’s instructions.

The pool of Riboseq human T-cell libraries was incubated with the Cas9-sgRNA complex for 2 hours at 37°C at molar ratios of 6.7uM sgRNA Pool:0.67uM Cas9:0.2pmol DNA Pool (1000 sgRNA:100 Cas9:1 DNA). Cas9 and the sgRNA pool were pre-incubated at 37°C for 15 min before addition of the DNA pool. Following the digestion, 15 U of RNase I was added to the sample and incubated at 37°C for 10 min. Cas9 was removed by treatment with 20 μg Proteinase K (5 mg/ml, Sigma Aldrich) for 15 min at 37°C, followed by heat-inactivation for 15 min at 95°C. The DNA was purified using the NucleoSpin Gel and PCR Clean-up Kit (Macherey-Nagel) and then subjected to 5 x PCR cycles using Illumina P5 (5’-AATGATACGGCGACCACC-3’) and P7 (5’-CAAGCAGAAGACGGCATA-3’) primers (purchased from IDT). The 100µl reaction was supplemented with 10 mM dNTPs, Phusion HF 5X Buffer and Phusion DNA Polymerase according to manufacturer’s instructions. The PCR product was purified using the Select-a-Size DNA Clean & Concentrator Kit using the ≥150bp (70µl) protocol, according to manufacturer’s instructions. The rRNA-depleted pool was prepared for deep sequencing on the Illumina NovaSeq System at GENEWIZ (Azenta Life Sciences, Germany).

**Table 1.**
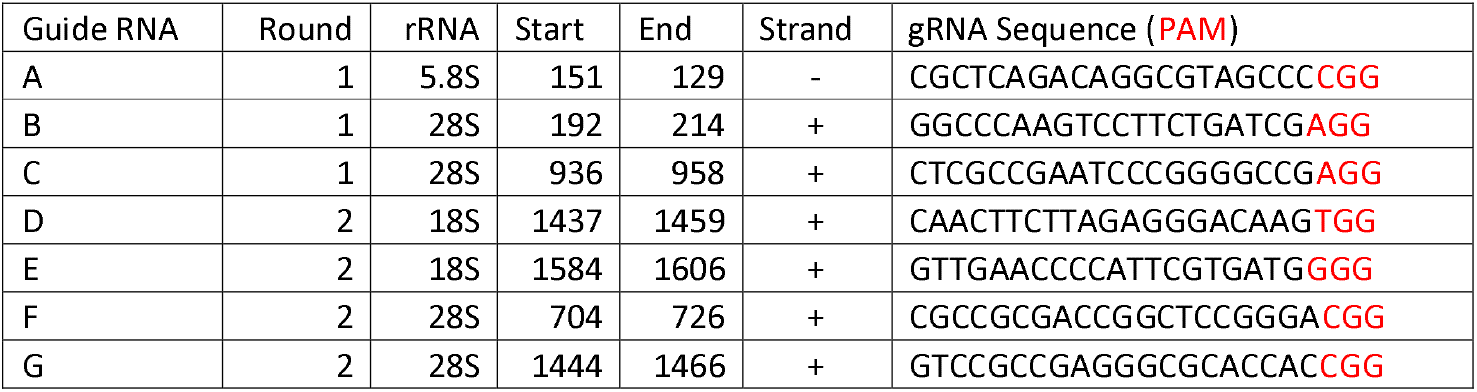
gRNA Sequence Design and PAM Sequences

**Table 2.**
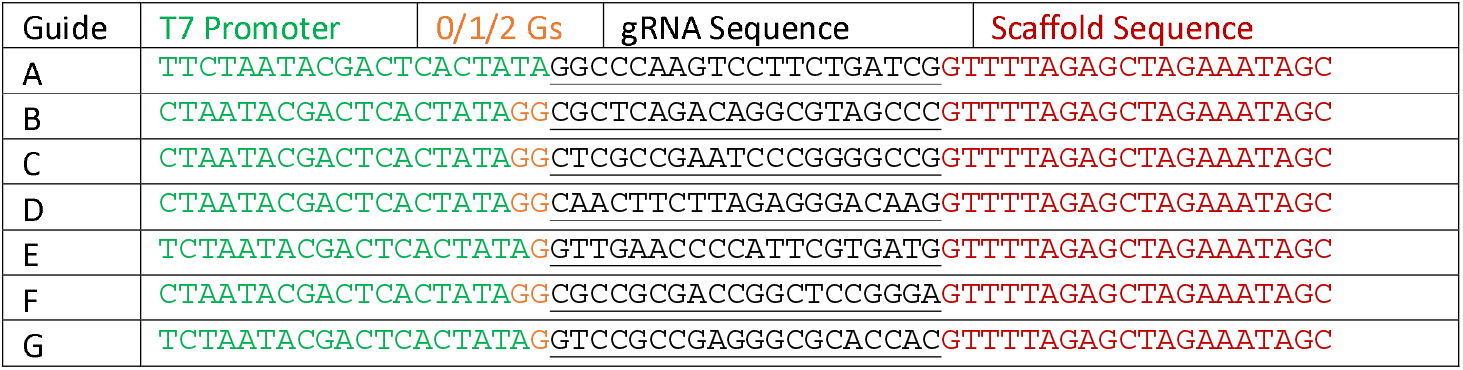
ssDNA Templates for rRNA Depletion

### Computational analysis of ribosome profiling and RNAseq data

Both stranded mRNAseq libraries and Riboseq libraries were sequenced 150PE on Illumina’s Nova-seq 6000 platform to depths of 20 million and 46-215 million raw read pairs per sample respectively.

For the Riboseq, the raw fastq files have the following sequence structure:

NNNNNNNNNNNN-QQQQ-rpf sequence - NNN – AGATCGGAAGAGCACACGTCTGAA

The first 12nt (N) are UMI additions. These are followed by (Q) a 4 nucleotide non-templated addition and the sequence of interest. The next 3 nucleotides (N) are an additional UMI. The adapter sequence is AGATCGGAAGAGCACACGTCTGAA.

Cutadapt (version 4.4) ^54^ was used to remove the linker sequence. Using EIRNABio’s proprietary script, the combined UMI and rpf sequence were used to collapse duplicates. (i.e. only one occurrence of each combined RPF and UMI sequence was retained). The UMIs (N) and non-templated addition (Q) sequence was removed from each read before further processing.

The following adapter sequences were removed from the RNAseq files using Cutadapt (version 4.4).

Read1 = AGATCGGAAGAGCACACGTCTGAACTCCAGTCA

Read2 = AGATCGGAAGAGCGTCGTGTAGGGAAAGAGTGT

To remove reads of artefactual origin, both the Riboseq and RNAseq were first aligned to rRNA, tRNA and snRNA/snoRNA using bowtie ^55^ (version 1.3.1 and settings bowtie -v 3). The reads were then aligned to the human transcriptome based on Gencode version 41 annotation ^56^. Genes annotated as pseudogenes, processed_transcripts and incomplete transcripts were filtered from the transcriptome. Reads were then mapped with bowtie (version 1.3.1 and settings -a -m 100 -l 25 -n 2 --norc) using a two-step approach. First reads were aligned to transcripts annotated as protein coding. Reads which failed to align to this were next aligned to the remaining transcripts annotated as non-coding. This approach reduces the number of reads which are ambiguously mapped thereby enabling better estimation of gene expression.

The transcriptome alignments were uploaded to EIRNABio’s proprietary bioinformatic analysis and visualization platform EIRNABio Connect (https://eirnabio.com/eirna-bio-connect/). Uniquely mapping reads are assigned a count of 1. Reads that map to >1 location are weighted according to the number of gene loci to which the read maps. For differential expression analysis, DEseq2 was used ^57^. Genomic alignments were visualized with JBrowse 2 ^58^. For the analysis of publicly available ribosome profiling data we used RiboSeq.Org resources ^59^.

## Acknowledgements

This work was supported by RSF grant no. 24-14-00213 (to D.E.A., data analysis), by RSF grant no 24-15-00332 (to Y.P.R., experiments with T cells). P.O’B was supported by Research Ireland Centre for Research Training in Genomics Data Science under grant number 18/CRT/6214. A.O’C was supported by Irish Research Council Enterprise Partnership Studentship Award EPSPG/2020/461. G.L. and P.V.B. were supported by Taighde Éireann – Research Ireland Frontiers for the Future Award (20/FFP-A/8929 to P.V.B.)

## Author contributions

D.E.A, Y.P.R., A.V.S., P.V.B. and G.L. designed experiments. A.V.S. and A.O’C. performed experiments. D.E.A., P.O’B., P.O’C., A.D.F. and A.M.M. analyzed and interpreted data. D.E.A. wrote the paper with input from all authors. All authors discussed the results and commented on the paper. All authors approved the final draft and agreed to the submission for publication.

