## Supplementary Figures for "The earliest changes in the translatome upon human T cell activation"

### Extended Data Figures

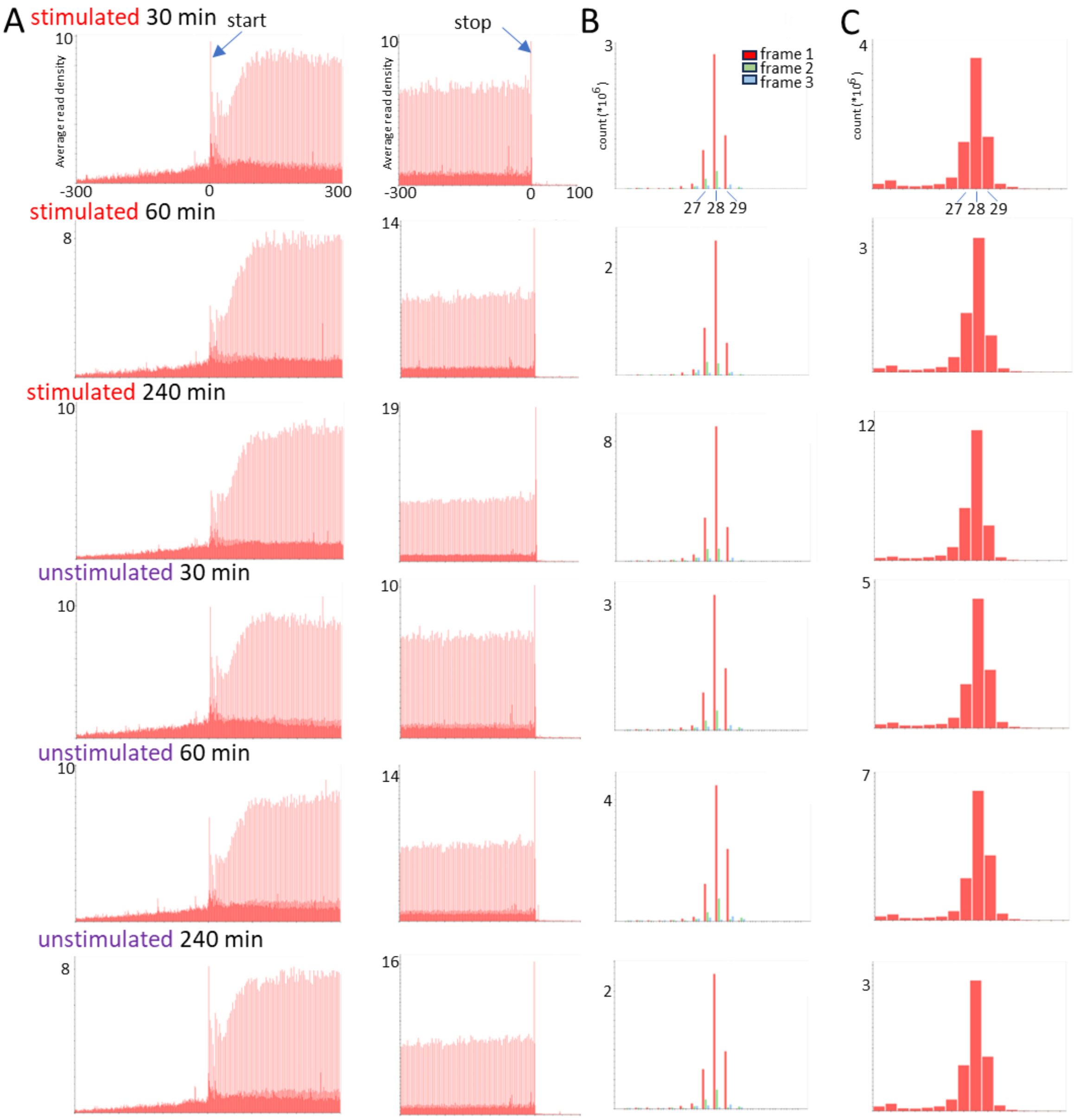


### Extended Data Fig. 1: Characteristics of Riboseq datasets

A) Metagene profiles, B) Triplet periodicity, C) Read length distribution

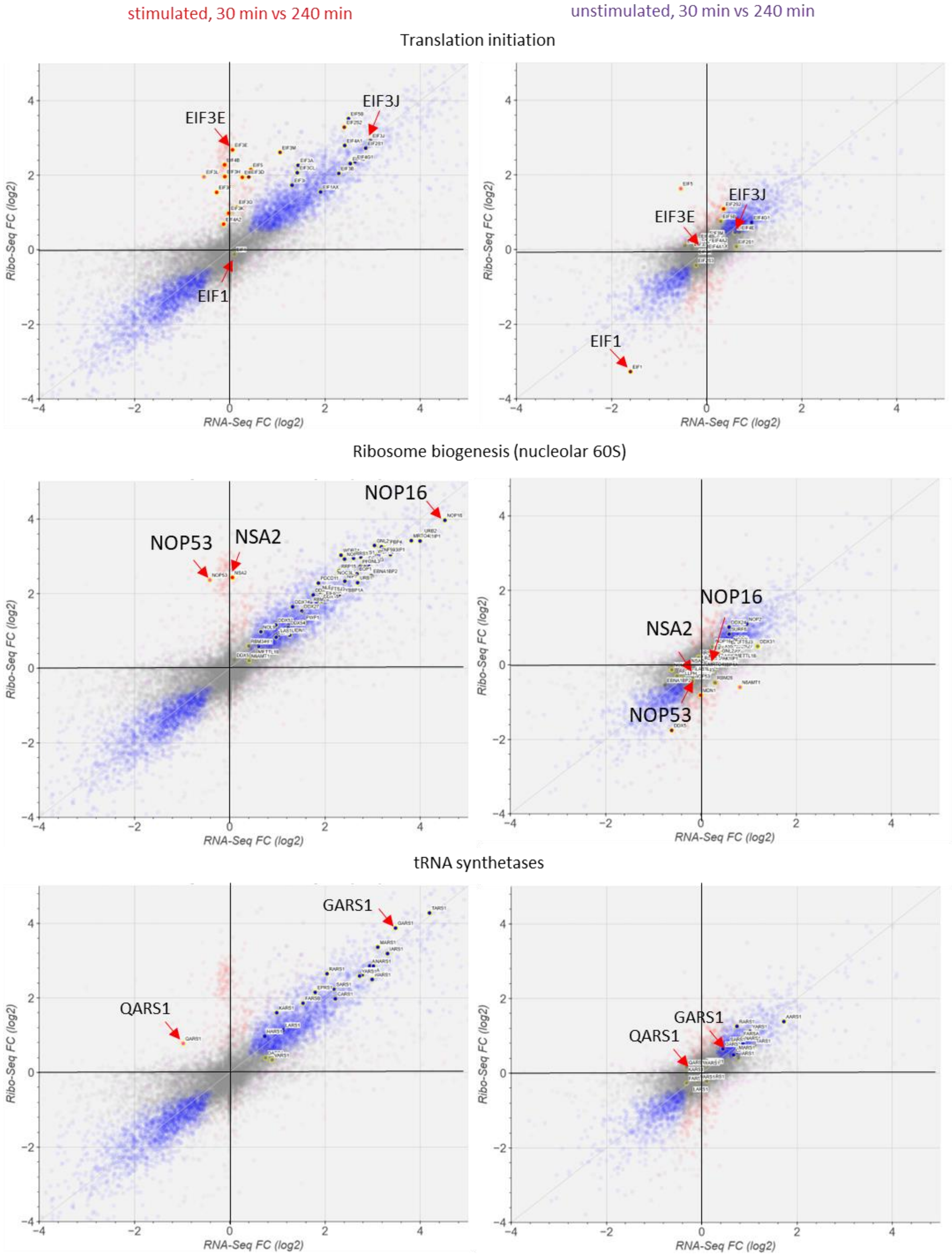


**Extended Data Fig. 2. Differential gene expression of selected genes of protein synthesis machinery**

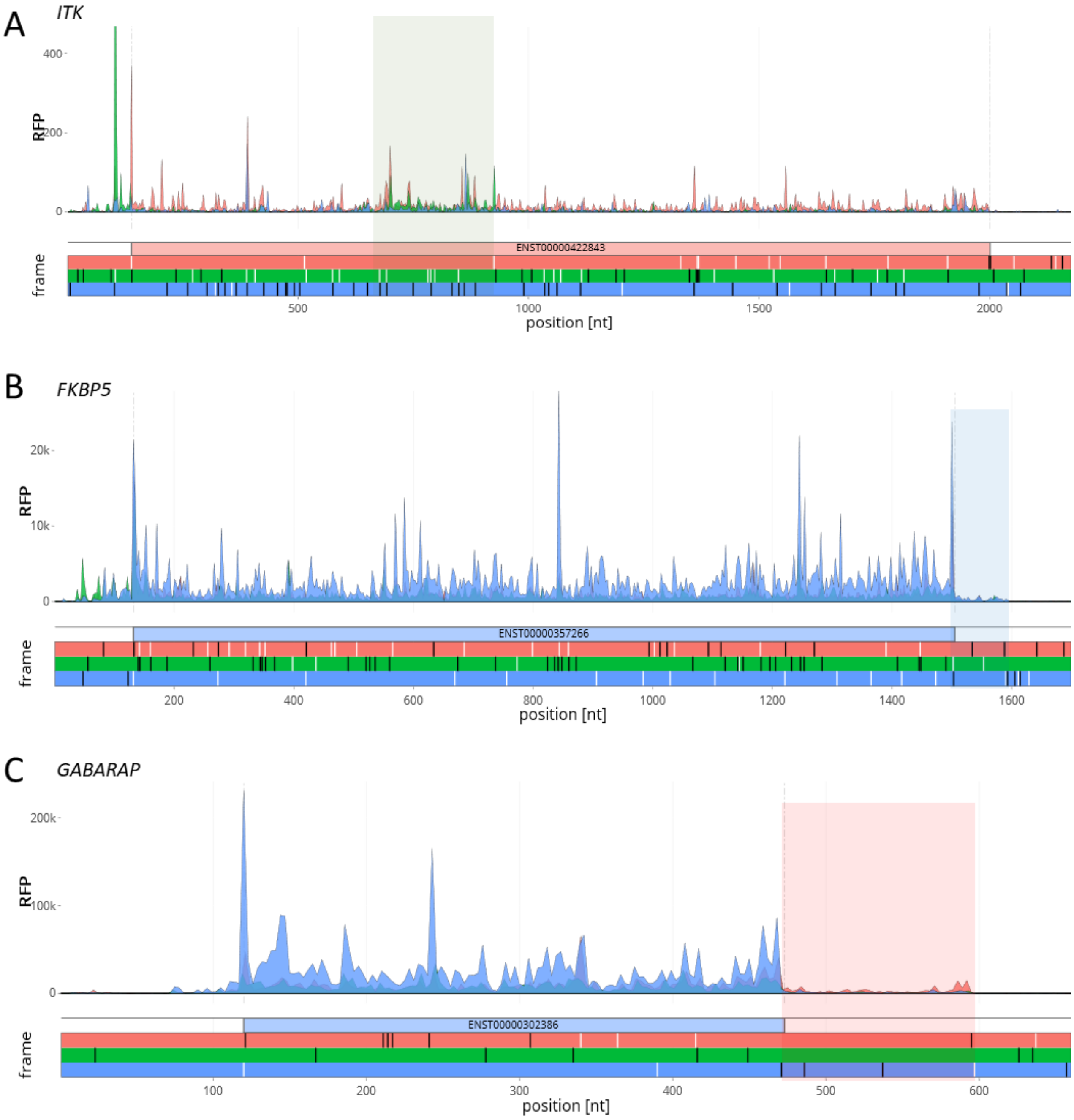


**Extended Data Fig. 3: Publicly available Riboseq profiles from Ribocrypt browser**

A) *ITK*, B) *FKBP5*, C) *GABARAP*. Regions of interests are highlighted in colored boxes.

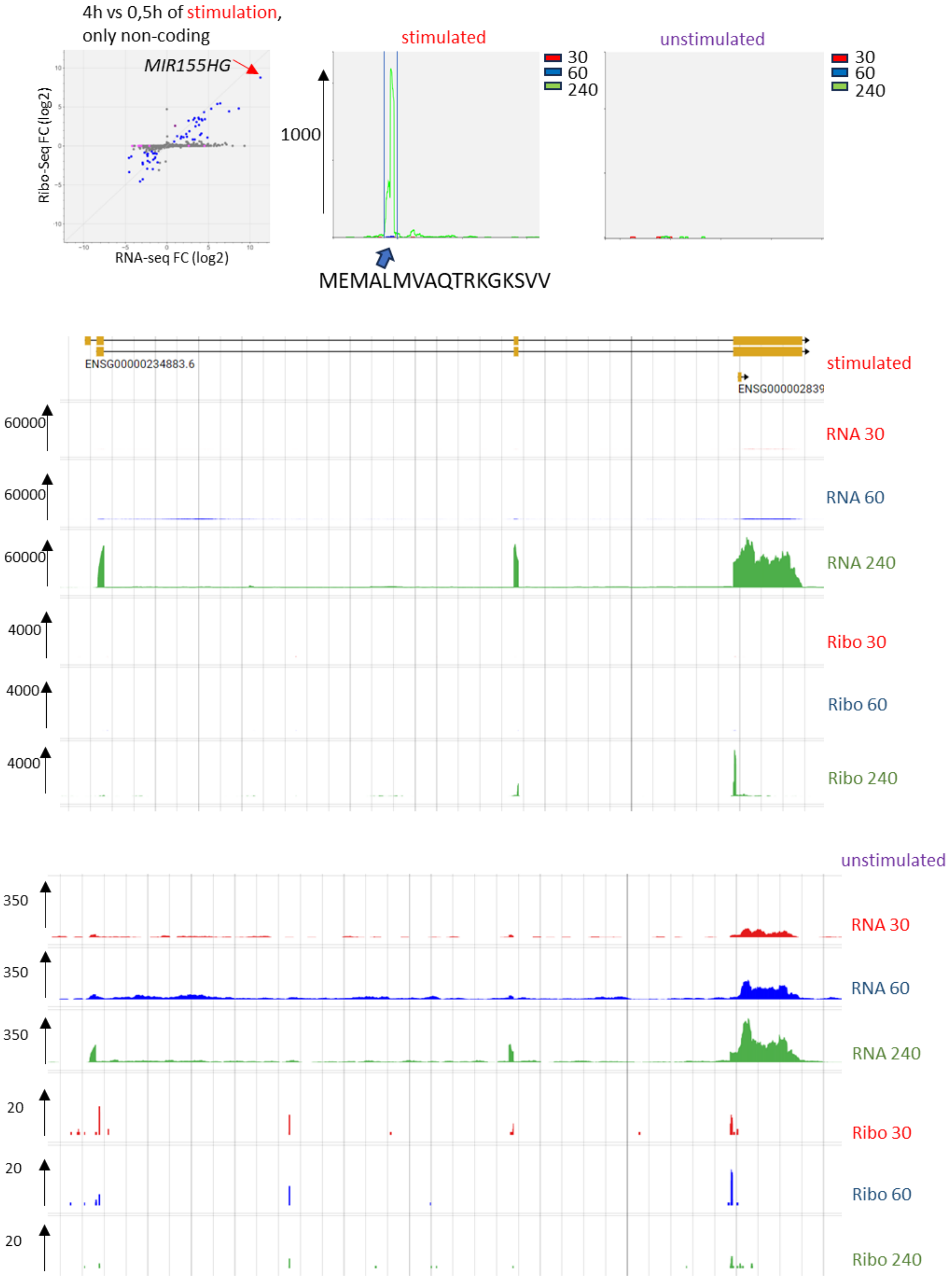


**Extended Data Fig. 4:** ***MIR155HG*** **expression in stimulated and unstimulated cells**

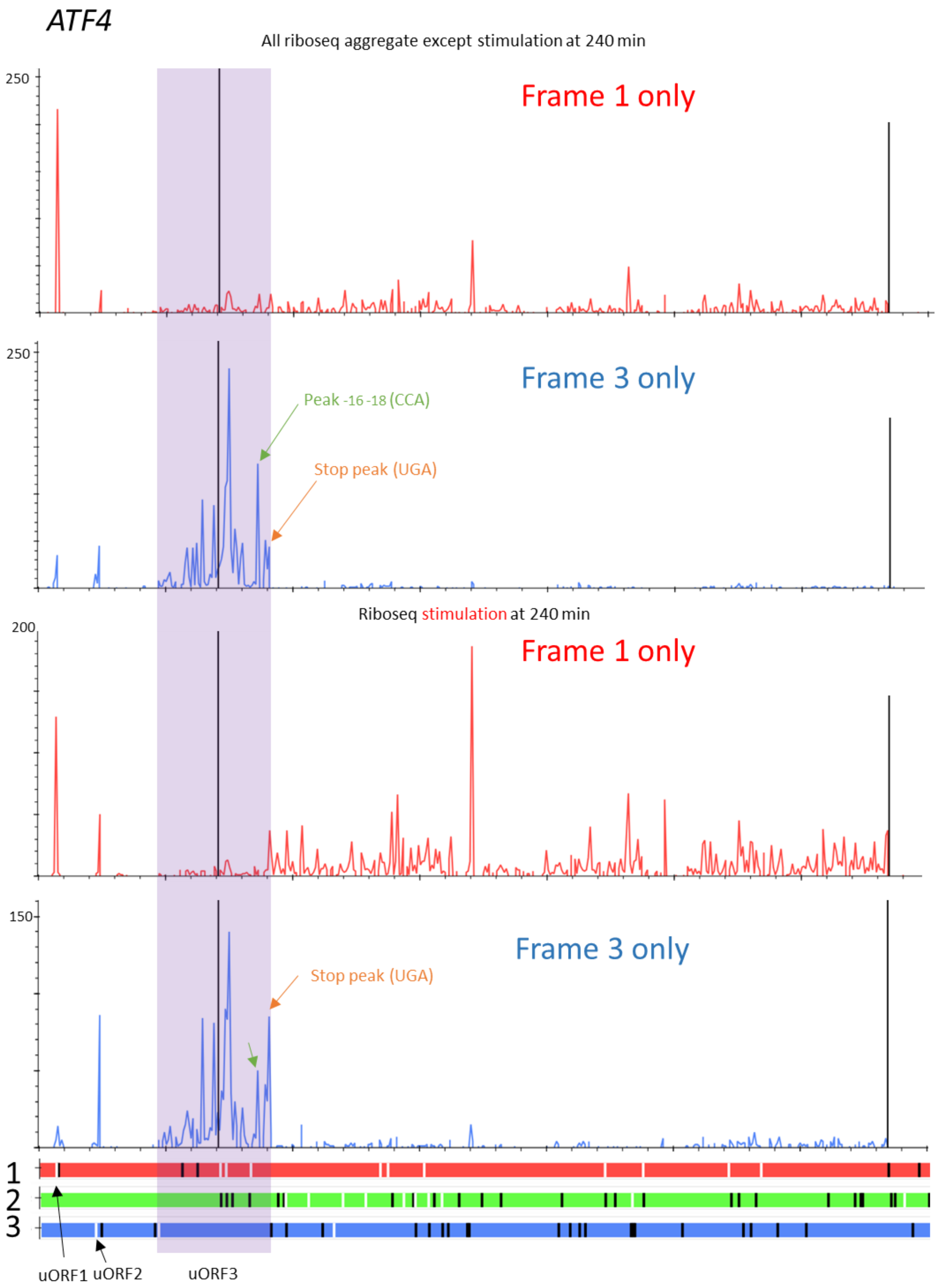


**Extended Data Fig. 5: Translation of *ATF4***

Subcodon profiles for *ATF4*. Only Riboseq footprints whose periodic distribution supports specific selected reading frames are shown. Selected peaks are highlighted with arrows.

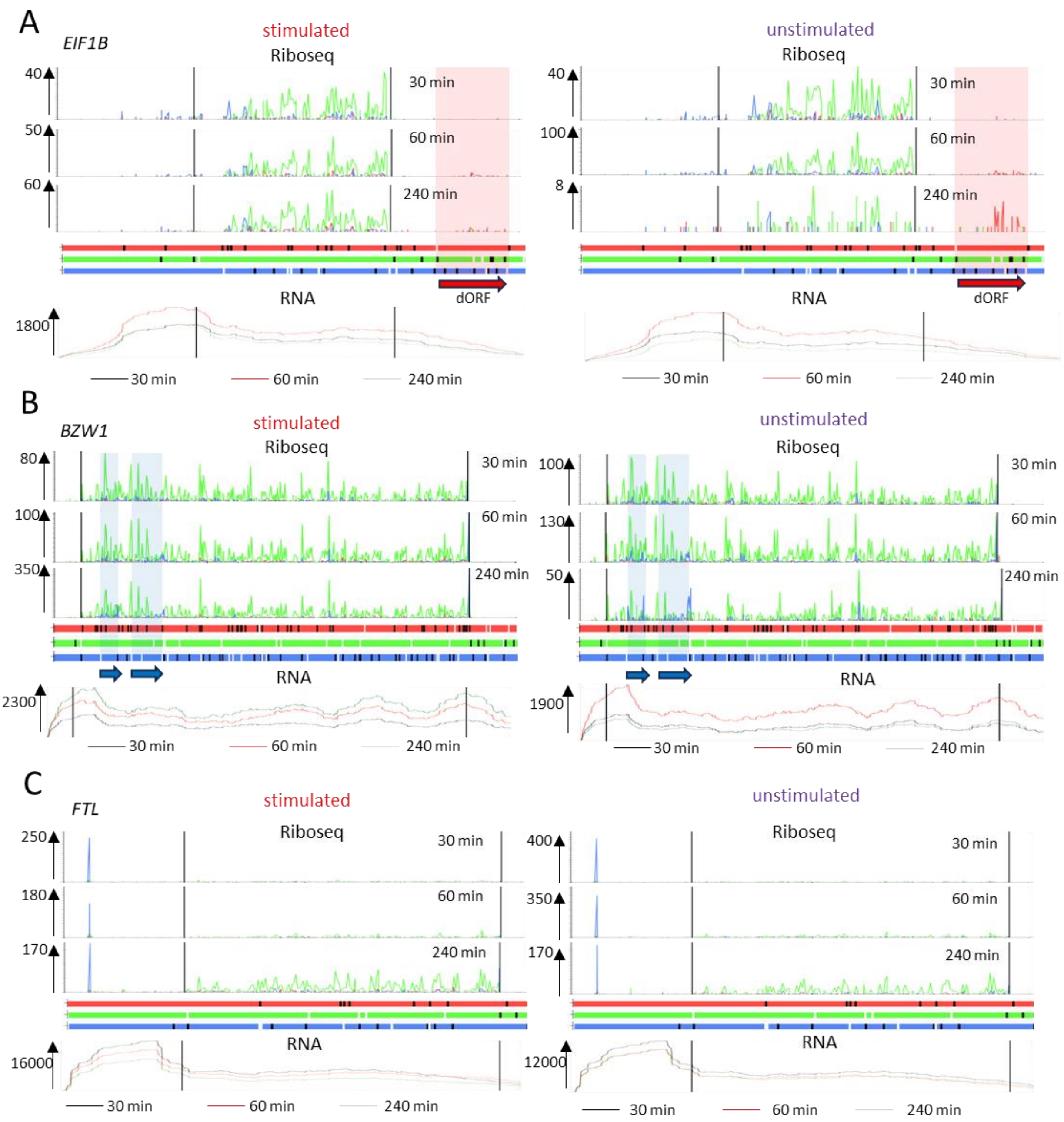


### Extended Data Fig. 6: Riboseq profiles of *EIF1B*, *BZW1* and *FTL*

Riboseq footprints are color-coded to match supported reading frames in ORF plot below. The numbers with arrows indicate the Riboseq scale on the y-axis. The region where Riboseq density changes is highlighted.
